# *Campylobacter* infection of young children in Colombia and its impact on the gastrointestinal environment

**DOI:** 10.1101/2024.05.06.592725

**Authors:** Zachary M. Burcham, Jessie L. Tweedie, AE Farfán-García, Vikki G. Nolan, Dallas Donohoe, Oscar G. Gómez-Duarte, Jeremiah G. Johnson

**Author notes:** Corresponding author: (JJ).

## Abstract

*Campylobacter* infections are a leading cause of bacterial-derived gastroenteritis worldwide with particularly profound impacts on pediatric patients in low-and-middle income countries. It remains unclear how *Campylobacter* impacts these hosts, though it is becoming increasingly evident that it is a multifactorial process that depends on the host immune response, the gastrointestinal microbiota, various bacterial factors, and host nutritional status. Since these factors likely vary between adult and pediatric patients in different regions of the world, it is important that studies define these attributes in well characterized clinical cohorts in diverse settings. In this study, we analyzed the fecal microbiota and the metabolomic and micronutrient profiles of asymptomatic and symptomatic pediatric patients in Colombia that were either infected or uninfected with *Campylobacter* during a case-controlled study on acute diarrheal disease. Here, we report that the microbiome of *Campylobacter-*infected children only changed in their abundance of *Campylobacter* spp. despite the inclusion of children with or without diarrhea. In addition to increased *Campylobacter,* computational models were used to identify fecal metabolites that were associated with *Campylobacter* infection and found that glucose-6-phosphate and homovanillic acid were the strongest predictors of infection in these pediatric patients, which suggest that colonocyte metabolism are impacted during infection. Despite changes to the fecal metabolome, the concentrations of intestinal minerals and trace elements were not significantly impacted by *Campylobacter* infection, but were elevated in uninfected children with diarrhea.

**Importance:** Gastrointestinal infection with pathogenic *Campylobacter* species has long been recognized as a significant cause of human morbidity. Recently, it has been observed that pediatric populations in low-and-middle income countries are uniquely impacted by these organisms in that infected children can be persistently colonized, develop enteric dysfunction, and exhibit reduced development and growth. While the association of *Campylobacter* species with these long-term effects continues to emerge, the impact of infection on the gastrointestinal environment of these children remains uncharacterized. To address this knowledge gap, our group leveraged clinical samples collected during a previous study on gastrointestinal infections in pediatric patients to examine the fecal microbiota, metabolome, and micronutrient profiles of those infected with *Campylobacter* species, and found that the metabolome was impacted in a way that suggests gastrointestinal cell metabolism is affected during infection, which is some of the first data indicating how gastrointestinal health in these patients may be affected.

## Introduction

*Campylobacter* species are a leading cause of bacterial-derived gastroenteritis in the world with a projected 96 million annual infections (1). Much of this prevalence is due to *Campylobacter*’s ability to asymptomatically reside within the gastrointestinal tracts of various animals, including livestock, and persist in the environment following dissemination from these hosts. Due to these sources, most human infections result from the consumption of contaminated water or undercooked meat. *Campylobacter jejuni* is the most common cause of human infection and is attributed to 80-90% of cases, followed by *C. coli*, which is responsible for 10-20% of infections. Despite the prevalence of these organisms, other species cause human infection and disease, including *C. concisus, C. hyointestinalis, C. lari, C. rectus, C. showae,* and *C. upsaliensis* (2). Following ingestion of these pathogens, clinical manifestation varies, but often includes the development of inflammatory diarrhea that may contain blood and be accompanied by abdominal cramps and/or fever. While infection is typically self-limiting and resolves in an average of 6 days, several lingering inflammatory disorders have been observed following infection, including Guillain-Barré Syndrome, inflammatory bowel disease, and reactive arthritis (2, 3). Beyond these outcomes, less inflammatory diarrhea and/or the development of an asymptomatic carrier state have been observed, particularly in pediatric patients (4).

In low-and-middle income countries (LMICs), pediatric patients are more likely to encounter *Campylobacter* species early in life and experience repeated infection or asymptomatic persistence than their counterparts in high-income countries (HICs) (5). As a result of these and other gastrointestinal infections, children in LMICs may develop environmental enteric dysfunction (EED), a condition that is characterized by villous blunting, decreased intestinal integrity, and reduced nutrient absorption (6). Unfortunately, the outcomes of developing EED include malnutrition, decreased cognitive development, reduced immune development, and decreased growth in these pediatric patients (7). While the effects of EED on intestinal health and nutrient absorption are beginning to be better understood, the specific impact of *Campylobacter* infection on the gastrointestinal environment of pediatric patients in LMICs remains unknown. To examine these effects, pediatric fecal samples were collected during a case-controlled study on acute diarrheal disease in Colombian and children were categorized based on *Campylobacter* infection status and whether they experienced diarrhea (8). Of these cohorts, we identified sub-populations of uninfected children that presented with diarrhea and *Campylobacter-*infected children that were asymptomatic. These sub-populations allowed us to identify changes to the gastrointestinal environment of pediatric patients of an LMIC that specifically occurred as a result of *Campylobacter* infection.

## Methods

### Study site

As previously described (9), the study was conducted in the Bucaramanga, Colombia metropolitan area. Bucaramanga is the capital city of the Santander department of Colombia. This city includes four municipalities, a population of 525,000, and 90% availability of basic utilities including those for water, electricity, gas, telephone, and garbage collection frequency of three times a week. The coverage of pediatric hospital beds in this region is 0.48 per 1000 for children under 12 years of age. The under-5 mortality rate in Colombia was 5.26 per 100,000 children in 2010, with acute diarrheal disease (ADD) being the third leading cause of morbidity among children less than five years of age in Santander, Colombia.

### Subject recruitment

Samples were collected during a study designed as a prospective, matched-for-age, case-control study to determine the etiology of ADD in children from two weeks to 59 months of age in the Bucaramanga metropolitan area, Colombia. The ADD study was approved by the Institutional Review Board, Vanderbilt University School of Medicine (IRB number 130327). From August 2013 to December 2014 subjects were recruited from emergency, inpatient, and outpatient facilities of major medical institutions including Unidad Intermedia Materno Infantil Santa Teresita (UIMIST), Centro de Salud el Rosario, Fundación Oftalmológica de Santander Carlos Ardila Lulle (FOSCAL), Hospital Local del Norte, Clínica Materno Infantil San Luis, and Hospital San Juan de Dios de Floridablanca in the Bucaramanga metropolitan area, Colombia. The inclusion and exclusion criteria for this study are included in Table 1.

**Table 1.** Inclusion and exclusion criteria for participants.

|  |  |  |
| --- | --- | --- |
| **Inclusion** | Children less than 60 months of age |  |
|  | Child who resides within the metropolitan area of Bucaramanga, Colombia |  |
|  | Presence or absence of acute, moderate-to-severe diarrhea<sup>1</sup> and/or vomiting within the past 10 days. |  |
|  | Diarrhea is moderate to severe and must meet at least one of the following criteria: | (i) Sunken eyes, confirmed by parent/caretaker |
| | | (ii) Loss of skin turgor by skin pinch ($\leq 2$ s slow or $>2$ very slow) |
|  |  | (iii) Intravenous rehydration prescribed or administered |
|  |  | (iv) Dysentery (1 or more bloody stools) |
|  |  | (v) Evaluated in the emergency department or admitted to the hospital for diarrhea |
| **Exclusion** | Children older than 60 months of age |  |
|  | Children who reside outside of the metropolitan area of Bucaramanga, Colombia |  |
| | Presence of chronic diarrhea ($>10$ days) or other comorbid conditions such as Crohn's disease or ulcerative colitis | |

<sup>1</sup>The World Health Organization defines diarrhea as 3 or more episodes of loose or liquid stools within 24 hours.

After informed written consent was obtained, an interview questionnaire was administered to the subject’s parents or guardians and recorded in Spanish by trained clinical researchers at enrollment and two and six weeks after. Data collected includes demographics, medical history, epidemiological factors, socioeconomic factors, nutrition, education, immunization history, water sources, and housing. In some cases, information was obtained about clinical manifestations including diarrhea, vomiting, abdominal pain, and dehydration. Access to the metadata and fecal samples was approved by and conducted under the University of Tennessee Institutional Review Board (#IRB-17-03795-XM).

### Sample Collection and Detection of Gastrointestinal Pathogens

Stool samples were collected from children aged two weeks to 59 months on either the day of enrollment or up to one-week post-enrollment from August to December 2014. Samples were collected in disposable plastic containers and transported to the Laboratorio de Investigaciones Biomédicas y Biotecnológicas (LIBB) at the Universidad de Santander, Bucaramanga, Colombia within four hours. Once samples were examined for color, consistency, presence of blood and/or mucus, aliquots were made and stored at −80°C. The samples were tested for viral, parasitic, and bacterial infections, including adenovirus, astrovirus, norovirus, rotavirus, sapovirus, *Entamoeba histolytica, Giardia lamblia, Cryptosporidium* spp*., Campylobacter* spp.*, Salmonella* spp.*, Shigella* spp.*, Yersinia* spp., enteroaggregative *Escherichia coli*, diffuse-adhering *E. coli,* enteropathogenic *E. coli*, enteroinvasive *E. coli*, and enterotoxigenic *E. coli*.

*Campylobacter* was detected by isolating on Campylosel agar (Biomerieux) and confirmed using API CAMPY (Biomerieux) and quantitative PCR (qPCR) as previously described. Briefly, human stool samples were diluted 1:10 in DEPC-treated water (Life Technologies), vortexed, and centrifuged. DNA was extracted using a QIAamp DNA stool mini kit (Qiagen) and stored at −20°C. A qPCR assay for *C. jejuni* and *C. coli* were conducted on a CFX96 Touch™ Real-Time PCR System as previously described (Bio-Rad). Each well included a 25 µl reaction mixture with 1 µl of DNA, 12.5 µl of TaqMan® environmental master mix, 9 µl of nuclease free water, 1 µl of each primer: cadF-Forward (5’-CTGCTAAACCATAGAAATAAAATTTCTCAC-3’) and cadF-Reverse (5’-CTTTGAAGGTAATTTAGATATGGATAATCG-3’) at a final concentration of 0.4 µM, and 0.5 µl of cadF-Probe (5’-[HEX]-CATTTTGACGATTTTTGGCTTGA-[BHQ2]-3’) at a final concentration of 0.2 µM. The cycling conditions were as follows: 95°C for 10 min, followed by 45 cycles of 95°C for 15 s and 55°C for 1 min.

### Generation and Sequencing of 16S rRNA Amplicon Libraries

Total microbial DNA was isolated from weighed, archival feces using the DNeasy PowerSoil Kit (QIAGEN) following the manufacturer’s protocol. DNA was stored at −20°C until 16S rRNA gene amplification. Primers targeting the V3-4 region of the 16S rRNA gene were used to create a single amplicon of approximately 460 bp: 16S Amplicon PCR Forward Primer (5’-CCTACGGGNGGCWGCAG-3’) and 16S Amplicon PCR Reverse Primer (5’-GACTACHVGGGTATCTAATCC-3’) according to the Illumina 16S Metagenomic Sequencing Library Preparation Guide. Once the library was quantified and normalized, it was pooled and sequenced on an Illumina MiSeq using v3 600 cycle kit (2 x 300 bp).

### Extraction of Patient Fecal Metabolites

Metabolites were extracted from participant fecal samples by adding 0.650 mL of Metabolic Extraction Solvent (MES) (20:40:40 H_2_O:ACN:MeOH + 0.1 M Formic acid) to 0.1 gram of sample and mixing thoroughly before adding another 0.650 mL of MES and chilling at −20°C for 20 minutes. Samples were centrifuged (4°C at 13,500 rpm) and the supernatant was transferred to a fresh microcentrifuge tube and stored at 4°C. The resulting pellet was resuspended in 0.2 mL of MES and chilled at −20°C for 20 minutes before centrifugation and transfer of the supernatant to a fresh microcentrifuge tube. The samples were put on ice and transported to the Biological and Small Molecule Mass Spectrometry Core (BSMMSC) of the University of Tennessee where the supernatants were dried under nitrogen and the resulting pellets resuspended in 0.300 mL of milliQ H_2_O before being transferred to autosampler vials.

### UHPLC-MS/MS Analysis of Patient Fecal Metabolites

The fecal metabolomes were defined by the BSMMSC at the University of Tennessee across two runs in December 2017 and March 2018. The samples above were separated on a Phenomonex Synergi Hydro RP, 2.5 um, 100 mm x 2.0 mm column at 25°C. Mobile phase elution of metabolites were: A) 97:3 Methanol to Water with 11 mM Tributylamine and 15 mM Acetic acid and B) 100% Methanol. The gradient used for mobile phase A during the 25-minute method with a flow rate of 0.2 mL/minute was 100% at minute zero, 80% at minute 5, 45% at minute 13, 5% at minute 15.5, 100% at minute 19 and 25, while the mobile phase B gradient equalized the percentage. The Exactive Plus Orbitrap used an electrospray ionization (ESI) probe operating in negative mode with a scan range of 72-1000 m/z.

### Inductively-coupled Plasma Mass Spectrometry (ICP-MS) Analysis of Fecal Micronutrients

The ICP-MS analysis was performed by the Spectroscopy and Biophysics Core of the University of Nebraska, Lincoln. The fecal samples were weighed and suspended in 1 mL of nitric acid (70% w/v). After overnight digestion at 65°C, the tubes were cooled, loaded in triplicate into 96-well plates, and diluted 20-fold with a solution of 50 ppb Ga in 2% nitric acid. Counts for boron, calcium, cobalt, chromium, copper, iron, magnesium, manganese, molybdenum, phosphate, potassium, selenium, sodium, sulfur, and zinc were normalized using this internal standard and were converted to concentrations using an external calibration curve. The instrument setup and acquisition method for ICP-MS were conducted as previously described (10).

## Statistical Analyses

### Epidemiological analysis of patient data

Descriptive statistics were generated for all clinical and demographic variables. For continuous variables, the association with being a *Campylobacter* case was determined by t-test or the non-parametric equivalent, the Mann-Whitney U-test. Categorical variables were analyzed with chi-square tests, with the exception of Fisher’s exact tests were used when expected cell counts were less than five. Wilcox two-sample t-tests, odds ratios, 95% confidence intervals, and p-values were determined with significance inferred at a p-value < 0.05.

### Analysis of patient fecal microbiomes

FASTQ sequences were imported into QIIME2 v2023.9 (11). Primers were removed from the sequences using Cutadapt v4.4 (12). Sequences were quality filtered, trimmed, and denoised into amplicon sequence variants (ASVs) using DADA2 (13) with the following parameters: --p-trunc-len-f 259, -- p-trunc-len-r 235, --p-trim-left-f 0, and --p-trim-left-r 0. Weighted 16S rRNA gene classifiers are trained with weights that take into account the fact that not all species are equally likely to be observed. If a sample comes from any of the 14 habitat types tested by Kaehler et al. (2019) (14) then weighted classifiers give increased classification precision. This includes the human gut habitat, therefore, the pretrained weighted Silva 138 99% OTUs full-length sequence classifier was used to classify the representative sequences generated by DADA2 with sklearn (15). The SILVA 128 database was utilized for phylogenetic tree creation with SEPP fragment insertion (16). SEPP fragment insertion performs a phylogenetic placement technique explicitly designed for 16S rRNA data to obtain improved phylogeny trees. Microbial features were filtered out if they were assigned to mitochondria, chloroplast, or not of bacterial origin. Further, features were removed to reduce noise if present less than 10 times in the data set and/or not found in at least 2 samples. After filtering, the data set included 38 samples and 938 features totaling a frequency of 3,649,980 feature counts. The median frequency per sample was 87,873 (minimum = 25,302; maximum = 261,609).

A rarefaction curve analysis was performed by generating alpha diversity metrics at a minimal sampling depth of 1 sequence to a maximum sampling depth of 100,000 sequencing per sample in 21 steps over 10 iterations. This determined that rarefying to a depth of 25,000 feature counts per sample captures the majority of the alpha diversity signal within the dataset; therefore, this sampling depth was used for calculating diversity metrics. The QIIME2 core-metrics-phylogenetic plugin was used to calculate alpha diversity metrics: Faith’s phylogenetic diversity, Peilou’s evenness, Shannon’s diversity, and ASV richness. The QIIME2 core-metrics-phylogenetic plugin was used to calculate beta diversity metrics: unweighted and weighted UniFrac distances (17) using the generated SEPP fragmentation phylogenetic tree and visualized with principal coordinates analysis (PCoA).

Differences of alpha diversity metrics between cohort groups, infection status, and symptomatic status were statistically tested using Kruskal-Wallis H tests. Weighted and unweighted UniFrac distances were statistically compared between cohort groups, infection status, and symptomatic status (i.e., diarrhea presence) using the adonis package permutational multivariate analyses of variance (formula = “infection status + symptomatic status + cohort group”; permutations = 999) (18). The non-rarified ASV feature table was collapsed by taxonomic ranks: genus, family, order, class, and phylum. Differential abundance of ASVs and each taxonomic rank was tested between cohort groups using the analysis of compositions of microbiomes with bias correction (ANCOM-BC) plugin in QIIME2 with the –p-conserve flag as recommended for small sample sizes and *Campylobacter*-uninfected asymptomatic as the reference level to compare against (19).

### Analysis of patient fecal metabolomes

The metabolomics data was generated in two separate runs; therefore, to reduce batch effects, only metabolite annotations that were detected in both runs were kept. Metabolite abundances were normalized by sample weight then scaled with pareto scaling using the scaling() function in the R v4.2.1 library MetaboAnalyze v1.3.1(20). The resulting feature table was imported into QIIME2 v2023.9 (11). The QIIME2 diversity plugin was used to calculate alpha diversity metrics: Peilou’s evenness, Shannon’s diversity, and richness, followed by beta diversity metrics: Canberra and Aitchison distances. Beta diversity metrics were visualized with t-SNE (t-distributed Stochastic Neighbor Embedding) (21) using the QIIME2 diversity tsne plugin with default parameters and --p-random-state 42. Differences of alpha diversity metrics between cohort groups, infection status, and symptomatic status were statistically tested using Kruskal-Wallis H tests. Canberra and Aitchison distances were statistically compared between cohort groups, infection status, and symptomatic status (i.e., diarrhea presence) using the adonis package permutational multivariate analyses of variance (formula = “infection status + symptomatic status + cohort group”; permutations = 999) (18). To determine if metabolome profiles were predictive of infection or symptomatic status, the QIIME2 sample-classifier plugin was used to predicted infection status or symptomatic status based on metabolite abundances using a Random Forest classifier trained and tested with nested cross-validation with the following parameters: --p-cv 5, -- p-estimators 999, --p-parameter-tuning, and --p-random-state 42. Results were analyzed by confusion matrices, receiver operating characteristic curves, or ROC curves, area under the ROC curves (AUC), and normalized frequency heatmaps of the determined top 20 important features for prediction of patient status. The importance values represent the contribution of each feature to the model’s predictive performance where the sum of all feature importances equals 1.

### Correlation network for estimating microbe-metabolite interactions

Annotated metabolite abundance tables and phylum microbial abundance tables were normalized to relative abundances. The tables were merged based on sample ID to only keep samples with both microbial and metabolite data. Pearson correlation coefficients were computed between the relative abundances of phyla and metabolites. For visualization, a clustermap with dendrograms was generated to represent microbe-metabolite correlations using seaborn.clustermap(). The ’average’ method was employed for hierarchical clustering which calculates the average distance between pairs of linked observations. The ’correlation’ metric was used to quantify the relationship between the phyla and metabolites. This metric ranges from −1 (perfect inverse correlation) to 1 (perfect correlation). A Pearson correlation network was constructed to visually represent strong correlations between phyla and metabolites using networkx (https://pypi.org/project/networkx/). A threshold was applied to consider only correlations with an absolute correlation greater than 0.2 to minimize noise. Edge width was adjusted based on the strength of the correlation where stronger correlations are represented by wider edges. Edges were colored to represent the positive or negative direction of the correlation. Microbial nodes were colored by whether the node represents a microbial phylum or metabolite.

### Analysis of micronutrient profiles

Outliers were detected using the ROUT method at Q=1% and removed prior to statistical analysis of both minerals and trace elements. Normality was examined using a Shapiro-Wilk test and differences between the CIS, CIA, CUS, and CUA cohorts were identified using a Kruskal-Wallis test (GraphPad Prism Software, La Jolla California USA). Significance was inferred at a *p-*value < 0.05.

## Results

### Analysis of sociodemographic variables found that few significantly differed between cohorts

Fecal samples from children under the age of five years old from Bucaramanga, Colombia were screened for the presence of gastrointestinal pathogens, including *Campylobacter*. *Campylobacter*-positive samples yielded detectable amplicons by qPCR using genus-specific oligonucleotides, whereas products could not be detected in *Campylobacter-*negative samples using the same oligonucleotides. From these fecal samples, a set of 20 *Campylobacter*-positive and 20 *Campylobacter*-negative were further classified as being from symptomatic or asymptomatic volunteers, depending on whether they presented with diarrhea at collection. This was done to control for non-specific impacts of diarrhea on the variables analyzed below. As a result, the following analyses were performed on the groups that are referred to as *Campylobacter*-infected, symptomatic (CIS) (n=14), *Campylobacter*-infected, asymptomatic (CIA) (n=6), *Campylobacter-*uninfected, symptomatic (CUS) (n=8), and *Campylobacter-*uninfected, asymptomatic (CUA) (n=12). Analysis of the sociodemographic data for the four groups found that none of the variables were significantly different (**Table 2**).

**Table 2.** Sociodemographics of study cohorts.

| Variable | CIS (%) | CIA (%) | CUS (%) | CUA (%) | p-value<sup>1</sup> |
| --- | --- | --- | --- | --- | --- |
| **Gender** |  |  |  |  | 0.47 |
| Male | 5 (35) | 3 (50) | 2 (25) | 4 (33) |  |
| Female | 9 (65) | 3 (50) | 6 (75) | 8 (67) |  |
| **Race** |  |  |  |  | 0.87 |
| Mestizo | 9 (64) | 5 (83) | 2 (25) | 12 (100) |  |
| White | 5 (36) | 1 (17) | 6 (75) | 0 (0) |  |
| **Age (months)** |  |  |  |  | 0.71 |
| 0 to 12 | 3 (21) | 2 (33.3) | 3 (37.5) | 6 (50) |  |
| 12 to 24 | 9 (64) | 2 (33.3) | 2 (25) | 2 (16) |  |
| 24 to 60 | 2 (14) | 2 (33.3) | 3 (37.5) | 4 (34) |  |
| **Total Subjects** | **14** | **6** | **8** | **12** |  |

### Other bacterial and viral pathogens were detected among patients

Because we defined the samples based on the presence of *Campylobacter* spp., 100% of the samples within the infected cohorts were positive for *Campylobacter* spp. at the time of collection while the uninfected cohorts were uniformly negative or undetectable for *Campylobacter* spp. When examining for other gastrointestinal pathogens, multiple were detected within both *Campylobacter*-infected and uninfected cohorts, including those with and without symptoms. For example, 15% of samples isolated from uninfected cohorts were positive for other gastrointestinal pathogens, while 85% were negative (**Table 3**). Importantly, although these infections were observed in both cohorts, significant correlations between a specific pathogen or cohort were not identified, with the exception of *Campylobacter*.

**Table 3.**
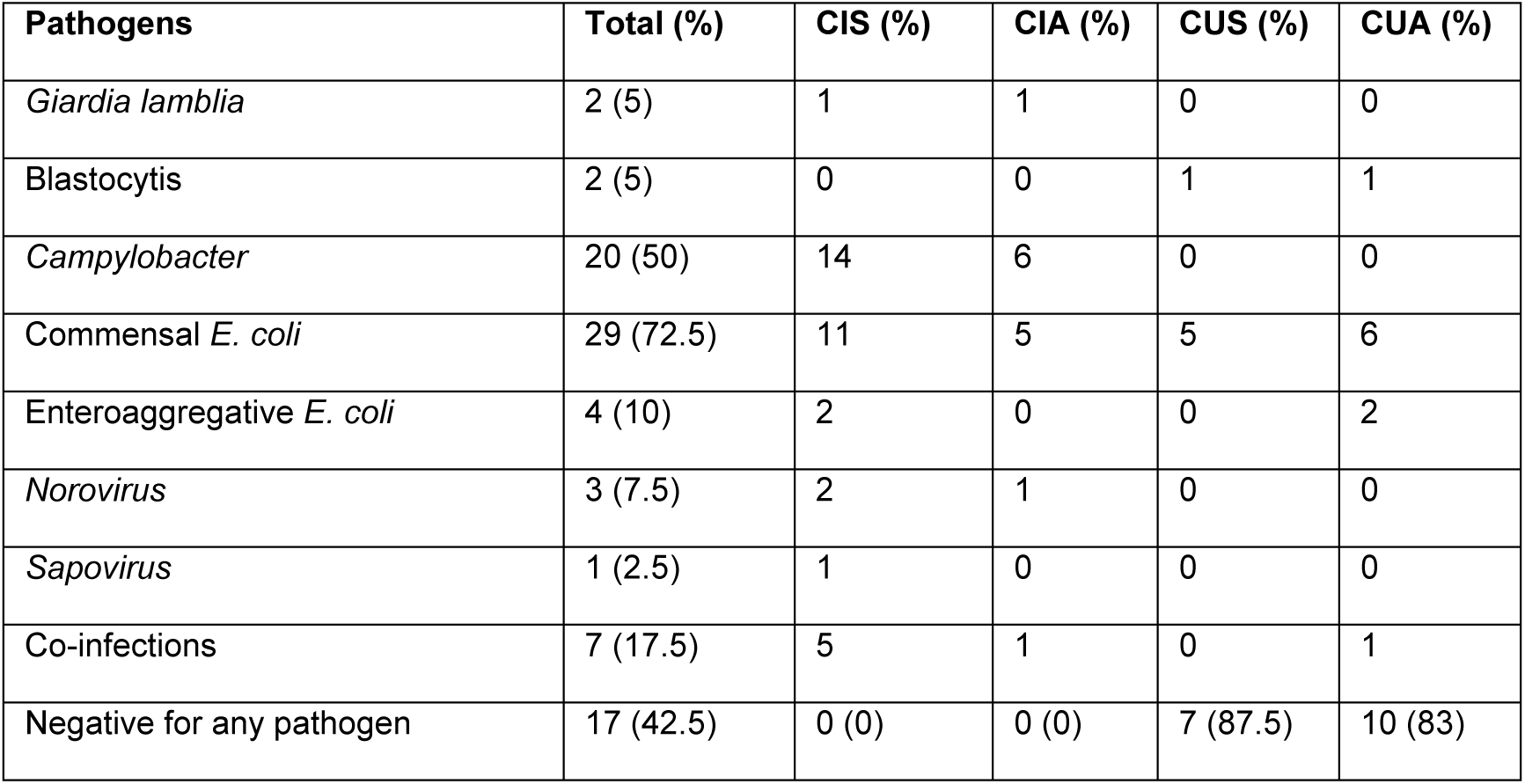
Co-infections detected among four groups.

### *Campylobacter* was detected in the microbiomes of all patient cohorts

Diversity analysis revealed no significant difference in ASV richness, Peilou’s evenness, Shannon’s diversity, and Faith’s phylogenetic diversity between the four cohorts (CIS, CIA, CUS, and CUA) (**Fig. S1**), *Campylobacter* infection status (**Fig. S2**), or if symptomatic (i.e., presence of diarrhea) (**Fig. S3, Table S1**). Similarly, weighted and unweighted UniFrac distances were not significantly impacted by cohort, *Campylobacter* infection status, or if symptomatic (**Fig. 1A; Fig. S4, Table S2**), suggesting neither the presence of *Campylobacter* nor the presence of symptoms systematically impacted patient microbial community composition.

**Figure 1.**
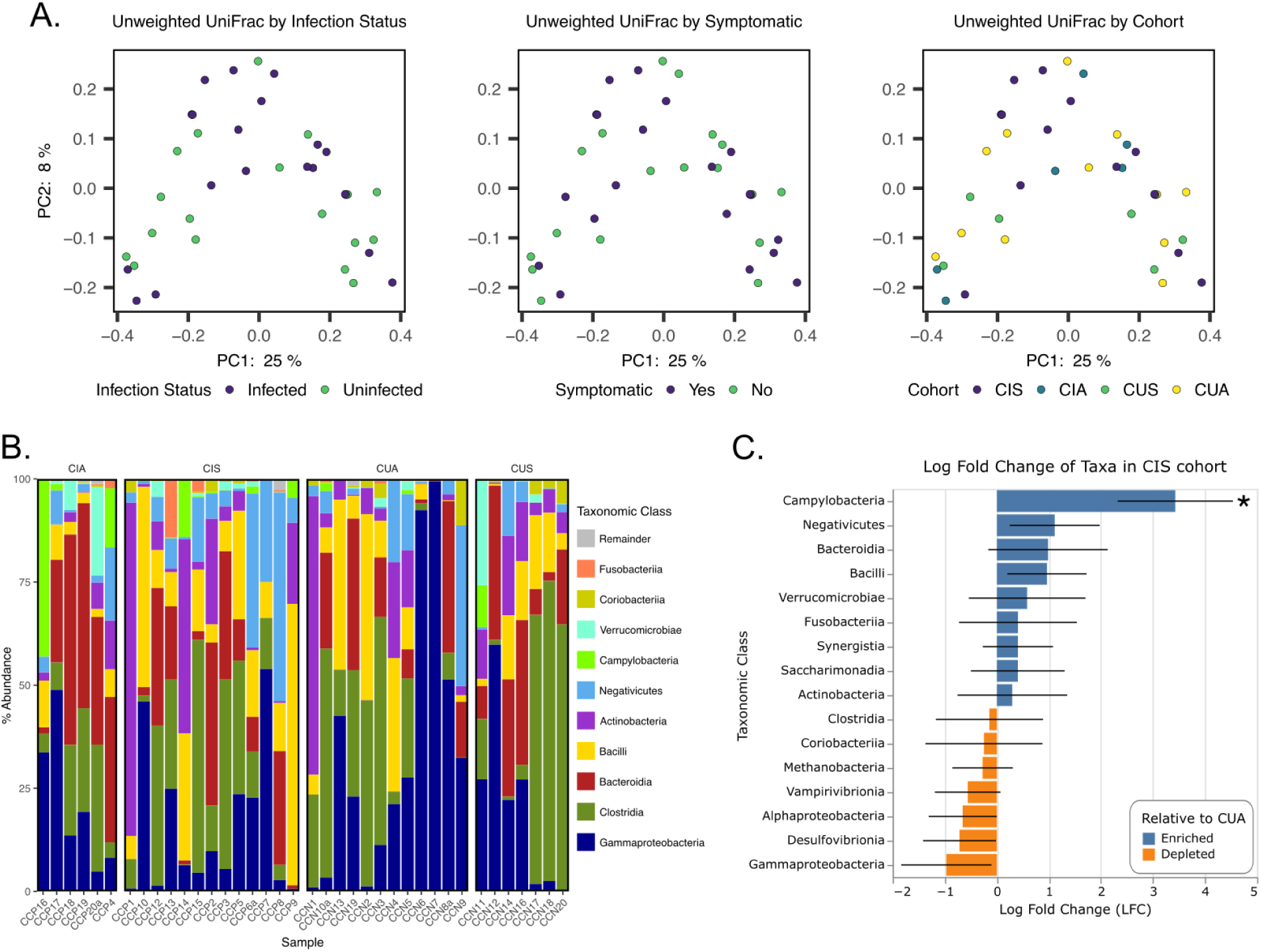
Microbiome analysis of patient cohorts. **A.**) Unweighted UniFrac distances visualized on PCoA plots to represent community similarity between infection status, symptomatic status, and cohort. **B.**) Relative abundances of the top 10 taxonomic classes detected in samples grouped by cohort. Classes outside of the top 10 relative abundance are collapsed as a remainder group. **C.**) The log fold change of taxonomic classes in the *Campylobacter*-infected, symptomatic cohort group relative to the uninfected, asymptomatic cohort group. Significance denoted by ‘*’ for FDR-corrected p-value < 0.05.

Exploration of cohort microbial community composition revealed that the most prominent classes across cohorts on average were Actinobacteria (10.6 ± 2.9%), Bacteroidia (16.7 ± 2.6%), Bacilli (13.7 ± 2.5%), Clostridia (21.5 ± 3.6%), Negativicutes (8 ± 2%), and Gammaproteobacteria (23.1 ± 4%) (**Fig. 1B**). Campylobacteria was detected in all cohort groups, even when *Campylobacter* was not detected in patients using the initial PCR, and was detected on average at 2.5 ± 1.2% of the community. It is important to note that in the previous study, testing for *Campylobacter* was specific for *C. jejuni* and *C. coli*, which could explain why *Campylobacter* was sequenced from all cohorts in our study, since non-*C. jejuni/C. coli* species would be identified using our methods. Despite the variation in average relative abundance of Campylobacteria between the cohorts and patients within each cohort, the designated *Campylobacter*-infected cohorts CIA and CIS had the highest average relative abundances of Campylobacteria at 9.8 ± 7% and 1.7 ± 1.1%, respectively. The Campylobacteria detected in the CIA cohort was largely dominated by two patients, CCP16 and CCP4, whose microbiomes constituted 43.1% and 14.3% Campylobacteria, respectively. The *Campylobacter*-uninfected, symptomatic (CUS) cohort had an average relative abundance of Campylobacteria similar to CIA at 1.5 ± 1.5%. The lowest average relative abundance of Campylobacteria was in the *Campylobacter*-uninfected, asymptomatic (CUA) at 0.25 ± 0.18%; however, this detection was largely driven by two patients, CCN5 and CCN10, and Campylobacteria was not detected in 8/12 CUA patients.

ANCOM-BC was performed to measure which taxonomic classes were enriched/depleted in each cohort and determine if these taxa were significantly differentially abundant in a cohort when compared to the CUA cohort. The only significant differential abundant class was Campylobacteria in the CIS cohort, which was 3.1 log-fold higher than in the CUA cohort as a whole (FDR-corrected p-value = 0.031) (**Fig. 1C; Table S3**). The CIA cohort was not significantly differentially abundant with Campylobacteria despite having the largest average Campylobacteria relative abundance (**Table S3**). This is likely due to the lack of consistent Campylobacteria detection between patients while Campylobacteria was more consistently detected in the CIS cohort. Taken together, these results suggests that while *Campylobacter* infection and diarrhea symptoms did not impact overall microbiome composition, Campylobacteria taxa were more prevalent in confirmed *Campylobacter*-infection cases and occasionally detected in non-infected and asymptomatic patients, suggesting a degree of low-level colonization across cohorts.

### Metabolomes are moderately predictive of *Campylobacter* infection in children

Since microbiome analysis by 16S rRNA gene characterization lacks the ability to give detailed information about the functional microbial activity, metabolomic analysis can give a glimpse of the intermediate phenotype mediating host–microbiome interactions (22). Using metabolomic analysis of the fecal samples by ultra-high-performance liquid chromatography tandem mass spectrometry (UHPLC–MS/MS), we observed 533 metabolites (peaks) of which 34 were assigned as specific metabolites in the database and were present in both metabolomic runs. Only the annotated metabolites present in both runs were analyzed as the metabolome profiles. Metabolome profiles had no significant difference in Peilou’s evenness, Shannon’s diversity, or richness between the four cohorts (CIS, CIA, CUS, and CUA), *Campylobacter* infection status, or if symptomatic (i.e., presence of diarrhea) (**Table S4**). Permutational multivariate analyses of variance revealed a significant difference in Canberra distances between symptomatic status but no other variables, while a significant difference between infection status was found in Aitchison distances, but no other variables (**Fig. S5; Table S5**). Due to how each distance metric is calculated, Canberra distances are often more sensitive to changes in low abundance features while Aitchison distances are often more sensitive to changes in more abundant features. These results suggest that changes in low abundance metabolites may be more indicative of symptomatic status than infection status, which might be more indicative of abundant metabolites.

To further explore the indicative metabolites, random forest machine learning classifier models were trained and tested via nested cross-validation to determine if the metabolite profiles were predictive of infection status or symptomatic status. We found that when predicting infection status, the true positive rate (i.e., sensitivity) was 0.737 and the true negative rate (i.e., specificity) was 0.6 for an overall accuracy of 66.7% (micro-average AUC = 0.71) suggesting metabolome profiles are moderately predictive of *Campylobacter* infection (**Table S6; Fig. 2A**). Two metabolites contributed to approximately 31% of the entire model’s predictive performance: glucose-6-phosphate (G6P) and homovanillic acid (HVA) (**Table S6**). Both of these metabolites are detected at higher frequency in *Campylobacter*-infected children. Using the same parameters to generate a model for predicting symptomatic status based on metabolome profiles, the sensitivity was 0.556 and the specificity was 0.619 for an overall accuracy of 60% (micro-average AUC = 0.66) suggesting metabolome profiles are not strongly predictive of symptomatic status (**Table S6; Figs 2B**). The feature importance was more evenly distributed in this model as the top two metabolites only contributed approximately 18% of the entire model’s predictive performance: homoserine/threonine and phenylalanine (**Table S6**). Both homoserine/threonine and phenylalanine were detected at higher frequency in symptomatic children.

**Figure 2.**
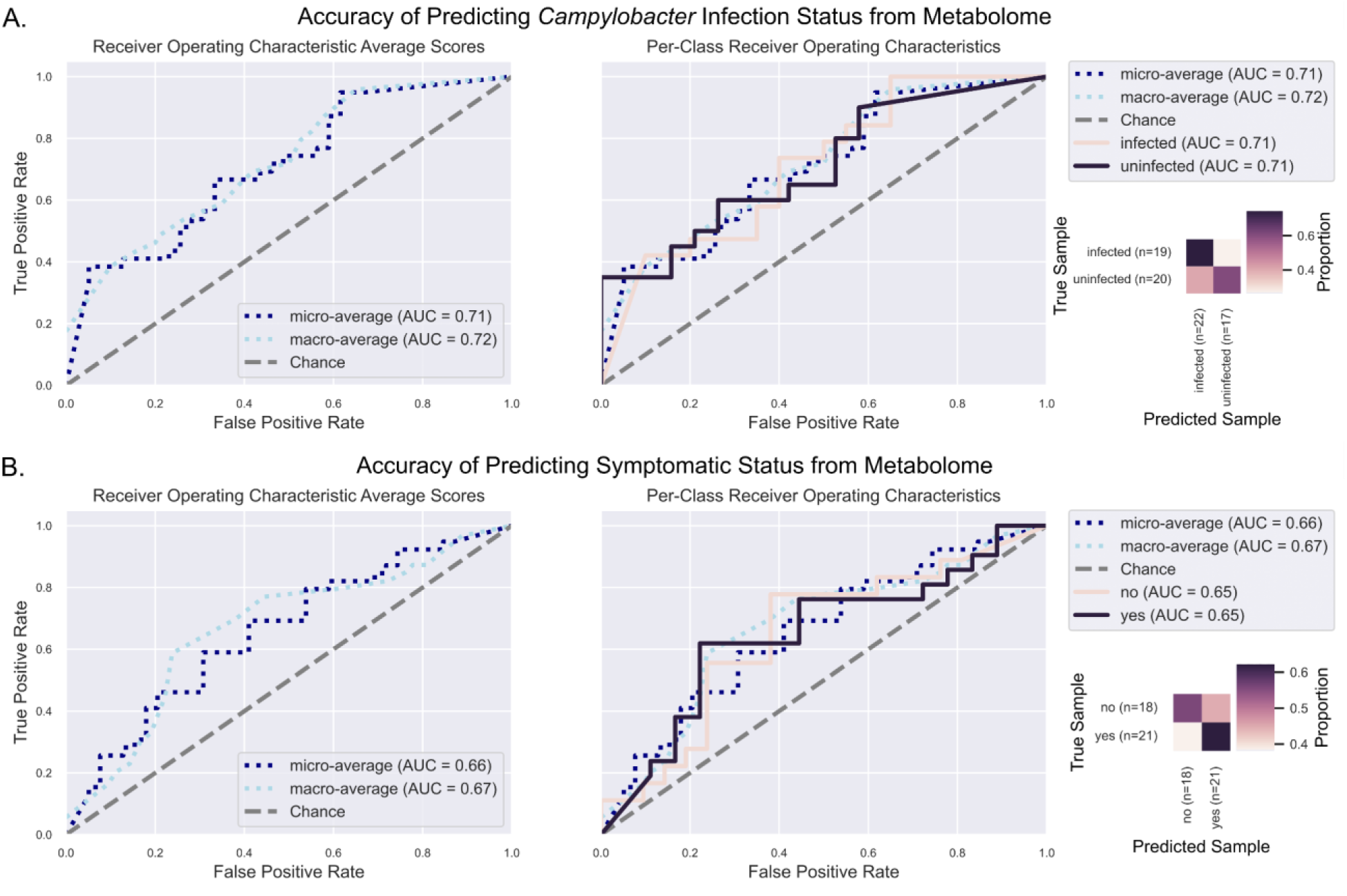
Random forest classifier with nested cross-validation accuracy results. The annotated metabolome profile abundances were used as features to predict **A.**) Campylobacter infection status and **B.**) symptomatic status for patients. Accuracy was assessed with receiver operating characteristic average scores and area under the curve (AUC) calculator coupled with a confusion matrix to visualize the proportion of true classifications predicted.

### Microbe-metabolite correlations highlight metabolites associated with Campilobacterota

Microbe-metabolite abundance correlations were used to explore potential interactions and co-occurrences of microbial phyla to the metabolites in the system regardless of infection status, symptomatic status, or cohort. We discovered that a complex correlation network exists between gut phyla and metabolites (**Fig. 3A-B; Table S7**). Campilobacterota was most positively correlated with malate (r = 0.345), taurine (r = 0.3), phenylalanine (r = 0.208), tyrosine (r = 0.198), and glucose-6-phosphate (r = 0.179), and most negatively correlated with succinate/methylmalonate (r = −0.209), lysine (r = −0.188), ornithine (r = −0.17), and serine (r = −0.167) (**Fig. 3A-B; Table S7**). The most important metabolites for predicting *Campylobacter* infection, glucose-6-phosphate and homovanillic acid, were most positively correlated with Fusobacteriota (r = 0.365) and Actinobacteria (r = 0.457), respectively (**Fig. 3A-B; Table S7**). Both of these phyla play a pivotal role in maintenance of gut homeostasis (23, 24); however, an unbalanced abundance of species within these phyla has been associated with pathological conditions including intestinal distress, gut microbiome dysbiosis, and colorectal cancers (25–28).

**Figure 3.**
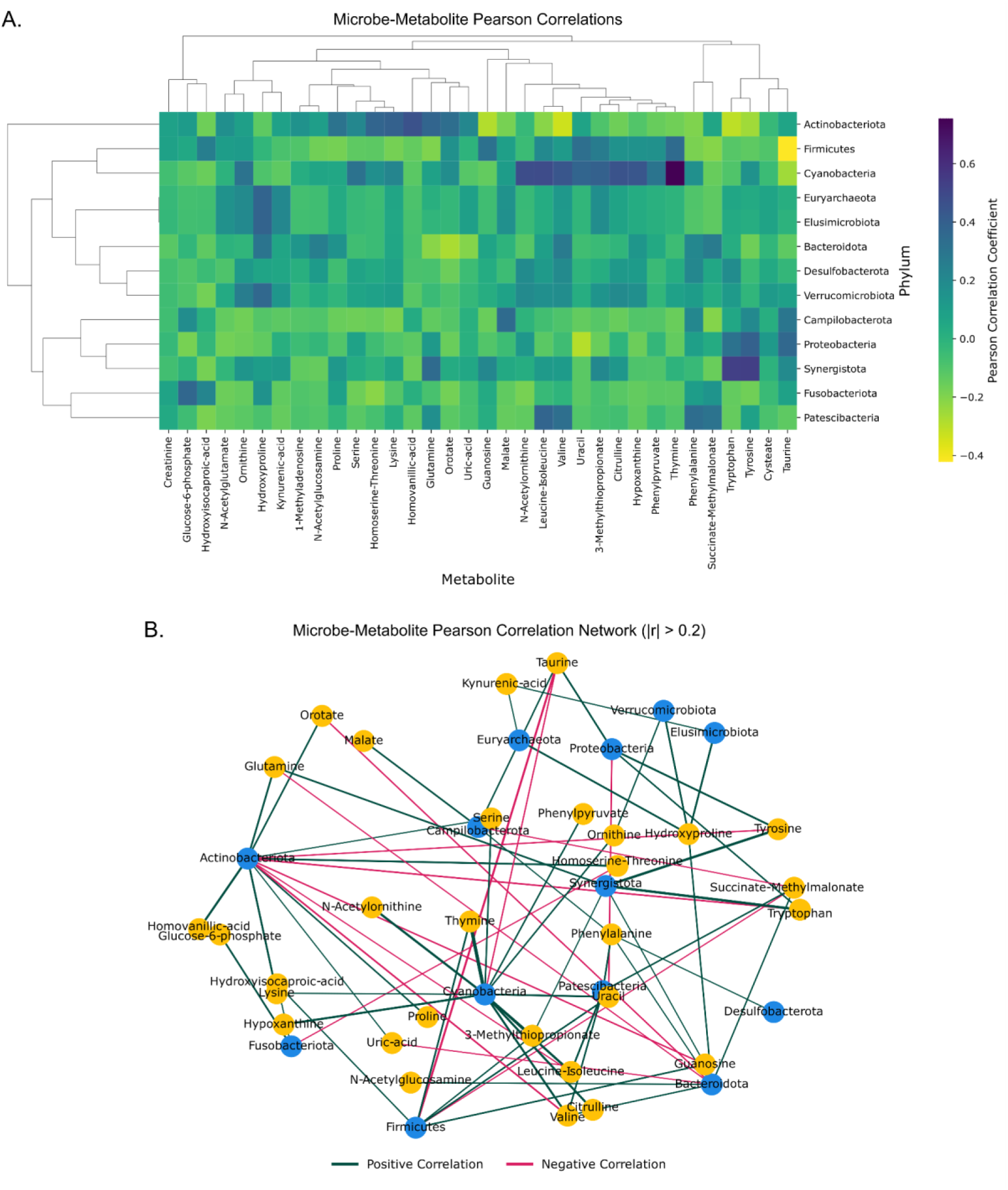
Microbe-metabolite abundance correlations. **A.**) Heatmap of all the Pearson correlations between phyla and annotated metabolites with dendrograms. **B.**) Network visualizing the top microbe-metabolite correlations (|r| > 0.2) across patients. Edge width was adjusted based on the strength of the correlation where stronger correlations are represented by wider edges. Edges were colored to represent the positive or negative direction of the correlation. Microbial nodes were colored by whether the node represents a microbial phylum or metabolite.

### *Campylobacter* infection does not lead to significant changes in fecal micronutrient concentrations

In addition to examining for changes to metabolites during pediatric *Campylobacter* infection, we also quantified the fecal concentrations of various minerals and trace elements. For minerals, we quantified the abundance of fecal calcium, iron, magnesium, phosphorus, potassium, sodium, sulfur, and zinc using inductively-coupled plasma mass spectrometry (ICP-MS) and normalized those results to the amount of fecal sample analyzed. Similarly, we determined the concentrations of the trace elements boron, cobalt, copper, chromium, manganese, molybdenum, and selenium. From this analysis, we found that whether *Campylobacter* infection was accompanied by symptoms or not (CIS and CIA), the mineral or trace element abundance of the fecal sample did not significantly differ from that of uninfected, asymptomatic children (CUA). This was observed for all elements examined with the exception of boron, which was significantly elevated in the feces of both CIS and CUS children. In contrast, children that were not infected with *Campylobacter*, but still experienced diarrhea (CUS), exhibited elevated levels of almost all minerals when compared to either asymptomatic cohort, including calcium, iron, magnesium, phosphorus, potassium, sulfur, and zinc. This trend was also observed for many of the trace elements, including boron, cobalt, copper, manganese, molybdenum, and selenium.

**Figure 4.**
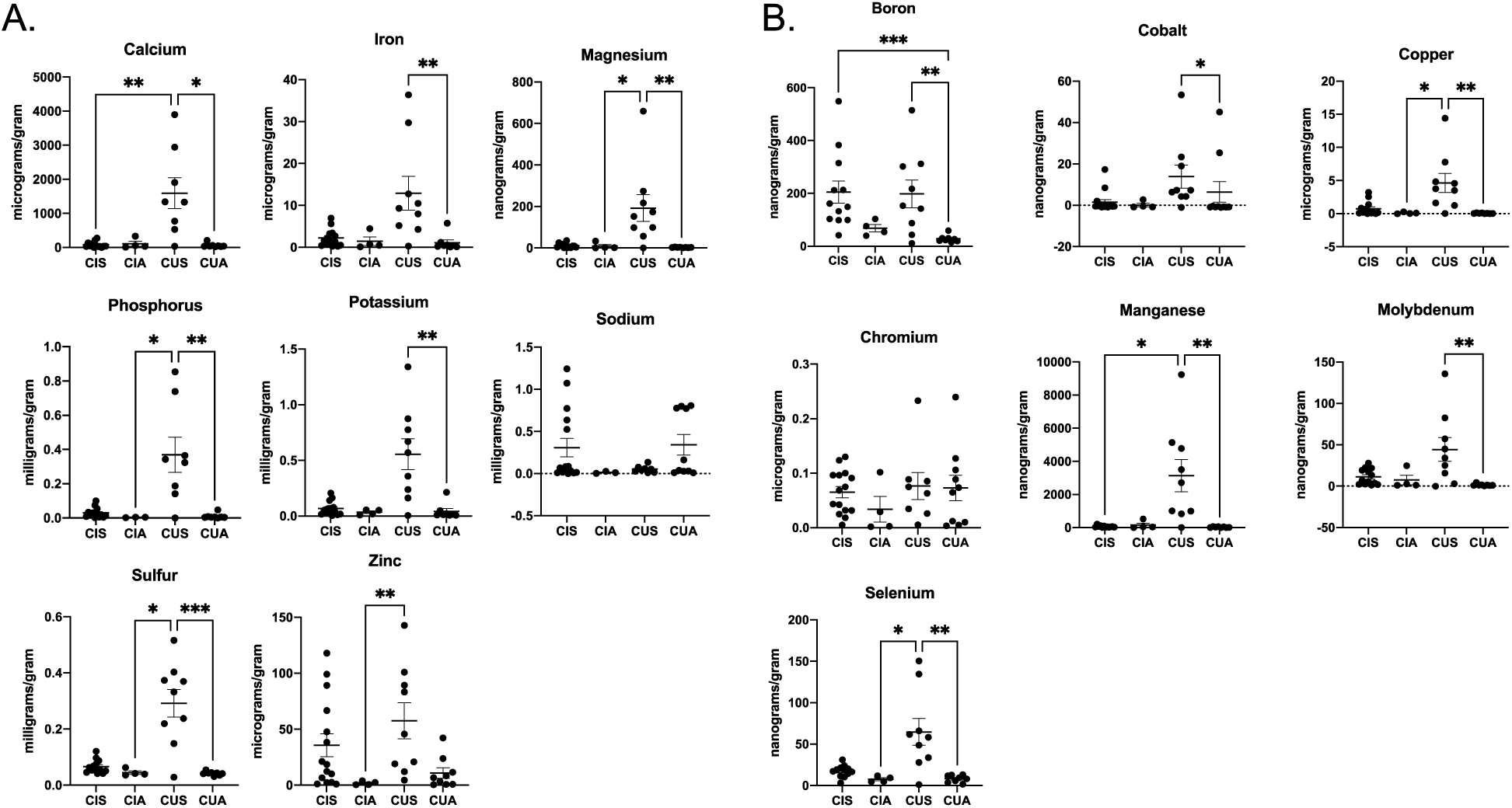
Mineral and trace element analysis of patient samples. Biologically-relevant minerals (A) and trace elements (B) were quantified for each patient sample by ICP-MS and concentrations were normalized by the weight of sample analyzed. Differences between CIS, CIA, CUS, and CUA cohorts were identified by one-way ANOVA. P-values <0.05*, 0.01**, and 0.001*** are shown.

## Discussion

*Campylobacter jejuni* is responsible for 96 million annual cases of bacterial-derived gastroenteritis worldwide, exceeding the number of cases produced by other well-known enteric pathogens including strains of pathogenic *E. coli* and *Salmonella* (2). To date, the impacts and mechanisms of *Campylobacter* infection in human hosts remain mostly unknown though it appears to affect patients in the developed and developing worlds differently. For example, *Campylobacter* infection has particularly profound impacts on pediatric subjects in the developing world where persistent colonization or re-infection leads to the development of EED, which can result in growth faltering and cognitive impairment (7). It was these effects on pediatric populations in the developing world that led us to examine changes to the microbiota, metabolome, and micronutrient abundances of Colombian children infected with *Campylobacter*.

The healthy human gut microbiota is dominated by anaerobes with dysbiosis occurring when the numbers of facultative anaerobic bacteria increase due to a reduction in epithelial hypoxia as a result of colonic inflammation (29). In this inflammatory environment, colonocyte metabolism shifts from mitochondrial β-oxidation of fatty acids toward anaerobic glycolysis, which results in high glucose and oxygen consumption and the release of lactate (29, 30). The resulting facultative anaerobic bloom, which is often dominated by members of the Enterobacteriaceae, leads to an altered nutritional environment through the generation of reactive oxygen species (ROS) and reactive nitrogen species (RNS) (31). While it is often recognized that ROS and RNS have antimicrobial activity, these molecules are also quickly converted to non-toxic compounds such as nitrate, which is a potent electron acceptor for the Enterobacteriaceae under anaerobic conditions (32). Further, ROS can facilitate the conversion of endogenous sulfur compounds into electron acceptors such as thiosulfate into tetrathionate (33). These environmental modifications would benefit *Campylobacter* since it is able to use both nitrate and tetrathionate for respiration (34–37).

Despite the effects of *Campylobacter* infection on pediatric patients in the developing world, its impact on the gastrointestinal microbiota is understudied. For example, a longitudinal study of young children from southern India found the gastrointestinal contents of these children were enriched for Campylobacterales, but the impact of colonization on the gastrointestinal microbial community was not investigated (38). In our study, we examined whether *Campylobacter* infection impacted the microbiota, but found the only significant change was the elevated relative abundance of the Campylobacteraceae, which increased from 2% in the uninfected cohort to 21% in the infected cohort at the phylum-level. Interestingly, we did not observe a change in Bacteroidetes or Firmicutes despite relatively high numbers of facultative anaerobes in both cohorts. This result suggests that children were already abundant for Enterobacteriaceae and when *Campylobacter* infection occurred, anaerobic members of the microbiota were already limited and no further decrease could be detected. Since *Campylobacters* are microaerophilic, with oxygen preferences of 2-12%, if epithelial hypoxia has already been lost in children enriched for Enterobacteriaceae, *Campylobacter* growth would be promoted and may allow for the persistent colonization that can occur in children from these communities (39, 40).

The two metabolites that contributed the greatest to the predicting the infection status of patient samples were G6P and HVA. It is interesting that G6P was predictive of infection status since characterized strains of *C. jejuni* are unable to use glucose as a carbon source due to the absence of several enzymes in the Embden-Meyerhof-Parnas glycolysis pathway, including a glucokinase which would be needed to generate G6P. As a result, the G6P detected in the infected patient samples is likely derived from either the microbiome or the host. In the case of the microbiome, since we did not observe significant changes in community member abundance during infection, the association of G6P with infection status could be caused by increased glycolysis of a microbe that is present in infected and uninfected cohorts. To examine this possibility, we performed a microbe-metabolite abundance correlation analysis and found that Fusobacteriota is connected to G6P abundance, which is intriguing since Fusobacteria were enriched, though not significantly, in CIS samples compared to CUA samples. Alternatively, the G6P associated with infection status could be derived from the host, which would suggest colonocyte metabolism shifted from short-chain fatty acid (SCFA) fermentation to glycolysis during pediatric infection. Importantly, a fermentation-to-glycolysis shift in colonocytes has been associated with promoting inflammatory signaling, differentiation to cancerous phenotypes, and decreased barrier function, all of which have been correlated with *Campylobacter* infection. HVA is a catecholamine metabolite that is predominantly recognized as a byproduct of dopamine degradation. Due to this association, most research involving HVA is focused on its differential abundance in cerebrospinal fluid, serum, and urine from patients with various neurological disorders, including alcoholism, depression, neuroblastomas, and parkinsonian syndromes (41–44). Because of this association, increased HVA in the feces of *Campylobacter-*infected patients could be sourced from the host due to the release of blood or serum into the intestinal lumen as a result of the immunopathology that is characteristic of campylobacteriosis. While we consider this the most likely explanation for the increased presence of HVA in the feces of these patients, we also considered whether a microbial component of the gut may be responsible for this. Using the same microbe-metabolite abundance correlation analysis above, we found that the phylum Actinobacteriota are associated with HVA, and that Actinobacteria were modestly, but non-significantly enriched in CIS samples when compared to CUA samples. It is unclear why Actinobacteria are correlated with HVA since there is little research on whether this organism can produce the metabolite, but it may be due to an apparent co-occurrence in gut-brain-axis studies where Actinobacteria and HVA are often associated with depression (9, 44). As a result, we consider it unlikely that gastrointestinal HVA is sourced from the microbiome.

Micronutrient analysis found that most minerals or trace elements were not significantly impacted during *Campylobacter* infection when compared to uninfected and/or asymptomatic cohorts, with the exception of boron, which also increased in children with diarrhea not infected with *Campylobacter*, suggesting that boron is generally elevated during gastrointestinal illness. Strikingly, children with diarrhea not infected with *Campylobacter* exhibited elevated fecal levels of almost all minerals and trace elements when compared to *Campylobacter-*infected and/or asymptomatic cohorts. It is unclear what is responsible for these changes, but initially examined the presence of diarrhea finding there were no significant differences between the symptomatic cohorts, but that the number of events and duration of diarrhea was significantly higher in *Campylobacter-*infected children (*p-*value < 0.03). Based on this finding, we suspect that symptomatic, *Campylobacter-*infected children lose minerals and trace elements at a rate that matches the absorption of those nutrients in asymptomatic children, which results in no apparent changes. In contrast, we hypothesize that symptomatic, *Campylobacter-*uninfected children developed gastrointestinal disease that resulted in elevated luminal micronutrients that were not lost at the same degree as *Campylobacter-*infected children. It would be interesting to determine whether this increased retention promotes reabsorption of nutrients and reduces the impact of illness on the host whereas nutrient loss during infection with *Campylobacter* results in less acquisition by the host, but this will need to be examined in the future using fecal and serum levels of these minerals and trace elements.

In conclusion, this study demonstrated that *Campylobacter* infection correlated with significant alterations to the gastrointestinal environment in pediatric populations. In the future, it would be interesting to conduct a similar, but longitudinal study to determine whether any of these children become persistently colonized and what the impact of that colonization is on the gastrointestinal environment. In addition, it would be desirable to obtain either blood or urine samples during such a study in order to more accurately determine what the systemic effects are of such an alteration. Such a longitudinal study would allow for the identification of children that exhibit decreased development and whether that outcome is potentially due to alteration in the gastrointestinal environment. These insights would contribute to EED prevention and/or treatment leading to enormously positive impacts on pediatric health, but also a substantially positive economic impact on the developing world.

## Data Availability

Microbiome and metabolome analysis files including code, intermediate files, and result files are publically-available on GitHub at https://github.com/BurchamLab/campy_colombia. Raw 16S rRNA FASTQ files are available on the NCBI Sequence Read Archive (SRA) under BioProject PRJNA1103705 and raw metabolomic files are available on the Metabolomics Workbench under ID 4784.

## Acknowledgements

The authors would like to thank Hector Castro-Gonzalez and Sara Howard from the University of Tennessee Biological and Small Molecule Mass Spectrometry Core for their help in preparing and analyzing samples for metabolomic analysis. We would also like to thank Joseph Jackson and Brittni Kelley at the University of Tennessee for providing support during the preparation of this manuscript.

## Funding

This work was funded from start-up funds provided to J.G.J. from the University of Tennessee as well as Seed Grant funding from the University of Tennessee Office of Research and Engagement’s (ORE) Strategic and Transformative Investments in Research (STIR) program to J.G.J. Also, this study was supported, in part, by NIH R01AI095346 to O.G.G.

### Author Contributions

ZB – data curation, formal analysis, investigation, methodology, validation, visualization, writing – original draft preparation, and writing – review & editing

JT – data curation, formal analysis, investigation, methodology, validation, visualization, writing – original draft preparation, and writing – review & editing

AFG – data curation, investigation, methodology, and writing – review & editing

VN – formal analysis, methodology, validation, and writing – review & editing

DD – methodology, supervision, and writing – review & editing

OG – conceptualization, methodology, project administration, resources, supervision, and writing – review & editing

JJ – conceptualization, funding acquisition, methodology, project administration, resources, supervision, and writing – review & editing

